# Single-nucleus RNA-Seq reveals a new type of brown adipocyte regulating thermogenesis

**DOI:** 10.1101/2020.01.20.890327

**Authors:** Wenfei Sun, Hua Dong, Miroslav Balaz, Michal Slyper, Eugene Drokhlyansky, Georgia Colleluori, Antonio Giordano, Zuzana Kovanicova, Patrik Stefanicka, Lianggong Ding, Gottfried Rudofsky, Jozef Ukropec, Saverio Cinti, Aviv Regev, Christian Wolfrum

**Author notes:** Correspondence (W.S.), (C.W.).

## Abstract

Adipose tissue usually is classified as either white, brown or beige/brite, based on whether it functions as an energy storage or thermogenic organ(Cannon and Nedergaard, 2004; Rosen and Spiegelman, 2014). It serves as an important regulator of systemic metabolism, exemplified by the fact that dysfunctional adipose tissue in obesity leads to a host of secondary metabolic complications such as diabetes, cardiovascular diseases and cancer(Hajer et al., 2008; Lauby-Secretan et al., 2016). In addition, adipose tissue is an important endocrine organ, which regulates the function of other metabolic tissues through paracrine and endocrine signals(Scheele and Wolfrum, 2019; Scherer, 2006). Work in recent years has demonstrated that tissue heterogeneity is an important factor regulating the functionality of various organs(Cao et al., 2017; Ginhoux et al., 2016; Park et al., 2018). Here we used single nucleus analysis in mice and men to deconvolute adipocyte heterogeneity. We are able to identify a novel subpopulation of adipocytes whose abundance is low in mice (2-8%) and which is increased under higher ambient temperatures. Interestingly, this population is abundant in humans who live close to thermoneutrality. We demonstrate that this novel adipocyte subtype functions as a paracrine cell regulating the activity of brown adipocytes through acetate-mediated regulation of thermogenesis. These findings could explain, why human brown adipose tissue is substantially less active than mouse tissue and targeting this pathway in humans might be utilized to restore thermogenic activity of this tissue.

## Main text

As one of the major endocrine tissues, the adipose organ is organized in different depots across the body and is composed of two different types: brown adipose tissue (BAT) and white adipose tissue (WAT). Each is comprised of a heterogeneous cell pool, which can be stratified into two groups. First, the functional cell pool of mature adipocytes, which represent 20 to 50% of the total cell content(Roh et al., 2018; Rosenwald et al., 2013), and second, the stromal vascular fraction (SVF), which includes preadipocytes, mesenchymal stem cells, fibroblasts, macrophages, immune cells, endothelial cells and vascular progenitors(Rosenwald and Wolfrum, 2014).

The main parenchymal cells of the adipose organ are adipocytes, which encompass three major cell types. White adipocytes store the energy taken up from the circulation into triacylglycerols. In contrast, brown adipocytes, and a less thermogenic efficient-related population referred to as beige or brite adipocytes, dissipate chemical energy in the form of heat to protect from cold temperature through non-shivering thermogenesis. This unique ability is enabled by the specific presence of uncoupling protein 1 (UCP1) in mitochondria(Jung et al., 2019). Notably, brown and beige/brite adipocytes share some morphological, biochemical and thermogenic characteristics, such as multiple small lipid droplets and a high abundance of mitochondria, packed with laminar cristae mitochondria, however, they also possess various distinct features. In rodents, classical brown adipocytes are located within distinct areas, like the interscapular BAT, whereas beige/brite cells are inducible and can be found within various WAT depots upon cold acclimation and β3-adrenergic receptor agonist stimulation(Rosenwald and Wolfrum, 2014). They arise from specific precursor cells, distinct from white adipocytes(Shinoda et al., 2015; Wu et al., 2012; Xue et al., 2015), but more recent data demonstrated a continuum in the expression pattern(Altshuler-Keylin et al., 2016; Shao et al., 2019). Additional studies identified four distinct human adipocyte subtypes differentially associated with thermogenesis and/or lipid storage(Min et al., 2019), and a recent study demonstrated the presence of brown adipocytes in mice with different thermogenic activities *in vivo*(Song et al., 2019), while another report showed immune cells and adipocyte interaction via *Il-10* using single-nucleus RNA-Seq (snRNA-seq)(Rajbhandari et al., 2019).

To comprehensively characterize the mature brown adipocyte subpopulations, we performed single-nucleus RNA seq(Grindberg et al., 2013; Habib et al., 2016, 2017) from interscapular brown adipose tissue (iBAT) of adult transgenic mice, in which red fluorescent protein (RFP) was expressed upon activation of the *Adipoq* promoter and localized in the nuclei(Straub et al., 2019) (**Fig. 1a, Fig. S1a**). Using RFP as a selection feature, we obtained and profiled 377 high-quality nuclei by full length scRNA-Seq with SMARTseq2(Picelli et al., 2014) (**Fig. S1a, Methods**), with 1,999 detected genes on average.

**Figure 1.**
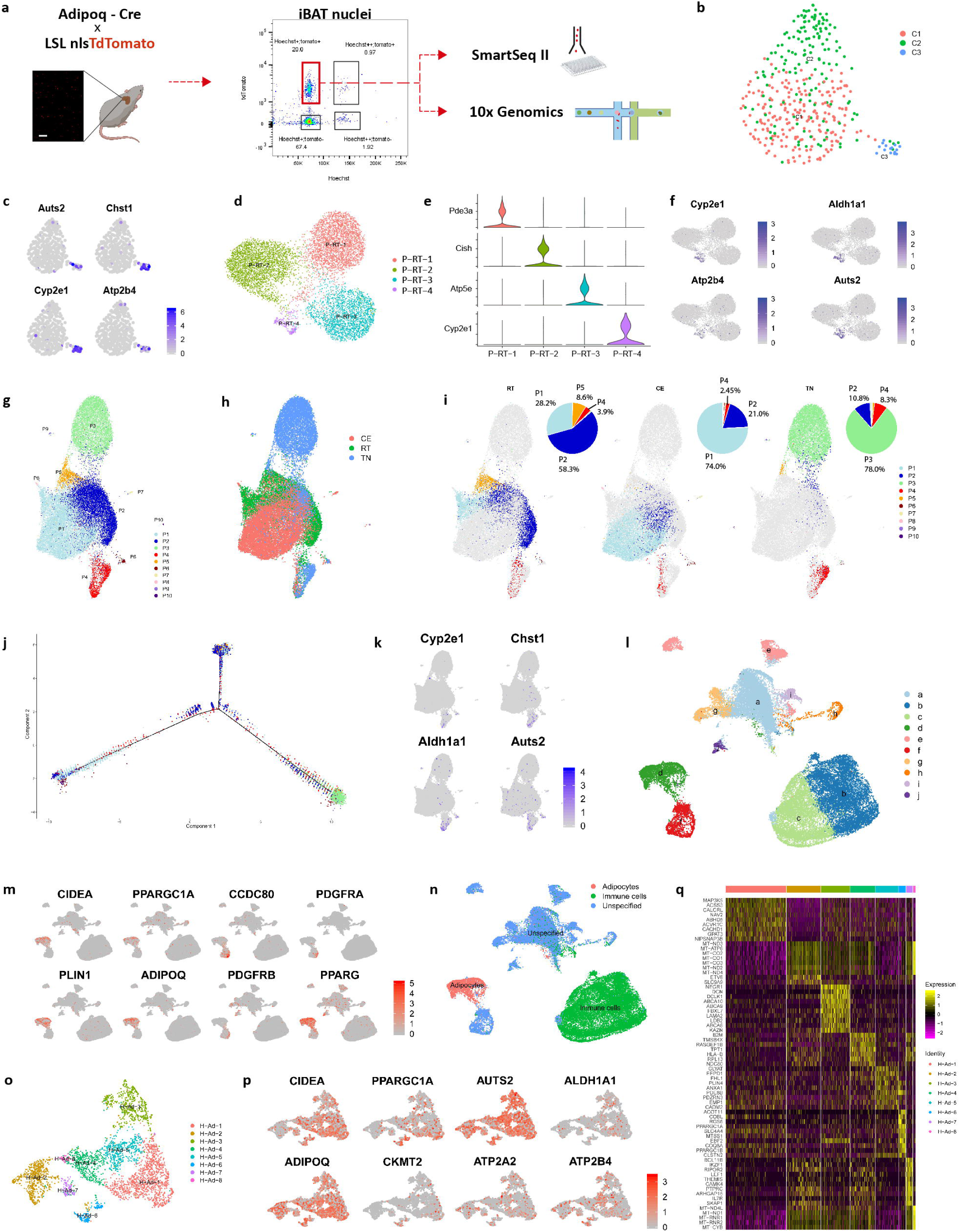
Identification of adipocytes populations in brown adipose tissue. (**a**) Schematic illustration of brown adipocyte nuclei sorting for single-nucleus RNA sequencing. (**b,c**) Single nucleus RNA sequencing of 377 adipocyte nuclei from brown tissue at RT, using the Smartseq2 protocol, yielding 1999 genes (medium) detected. (**b**) Unsupervised clustering shown as UMAP. (**c**) Feature plots for marker genes expressed in C3. (**d**-**f**) Single nucleus RNA sequencing of 8827 adipocyte nuclei from brown tissue at RT, using the 10x protocol. (**d**) Unsupervised clustering shown as UMAP. (**e**) Violin plot of *Pde3a, Cish, Atp5e, Cyp2e1* (**f**) Feature plots for marker genes expressed in P-RT-4. (**g**-**k**) Integrated single nucleus RNA sequencing of 28,771 adipocyte nuclei from brown tissue at TN, RT, CE, using the 10x protocol, yielding 1,265 genes (medium) detected. (**g**) Unsupervised clustering showed as UMAP. (**h**) UMAP plots grouped by different conditions. (**i**) UMAP plots split by different conditions. (**j**) Pseudotime plot of integrated adipocytes snRNA seq by monocle. (**k**) Feature plots for marker genes expressed in P4. (**l-q**) Single nucleus RNA sequencing of human deep-neck brown adipose tissue. (**l**) Unsupervised clustering of 42,295 nuclei from human brown adipose tissue, yielding 1,518 (medium) genes detected. (**m**) Feature plots for *CIDEA, PPARGC1A, CCDC80, PDGFRA*, PLIN1, *ADIPOQ, PDGFRB*, and *PPARG*. (**n**) Classification of adipocytes and immune cells by Garnett. (**o**) Adipocyte classification shown in UMAP. (**o**) Unsupervised clustering of 3,599 adipocyte nuclei from human brown adipose tissue, yielding 1,064genes (medium) detected. (**p**) Feature plot of *AUTS2, ALDH1A1, CIDEA, PPARGC1A, CKMT2, ATP2A2, ATP2B4 and ADIPOQ*. (**q**) Heatmap of the marker genes of each population of human adipocytes. Scale bar is 50 μm.

Unsupervised clustering of the profiles identified three adipocyte subsets, referred to as groups (C1, C2 and C3) (**Fig. 1b, Methods**). Canonical adipocyte markers *Adipoq, Plin1, Lipe, Cidec* and brown adipocyte markers *Ucp1, Cidea, Ppargc1a, Syne2*(Shinoda et al., 2015) were expressed in all groups (**Fig. S1b**), albeit at varying levels. Preadipocyte markers, such as *Cd34 and Ly6a (Sca1)* (**Fig. S1c**) were found in few cells in cluster C2, which could denote differentiating precursor cells which have a high adipogenic signature(Merrick et al., 2019; Schwalie et al., 2018). Trajectory analysis by RNA velocity and monocle suggested differentiation of these cells from population C2 to C1 (**Fig. S1d,e**). C3 cells (15 out of 377, blue) are a noticeably distinct group of brown adipocytes with distinctive marker genes, including *Cyp2e1, Chst1, Auts2* and *Atp2b4* (**Fig. 1c** and **Extended Data Table 1**). We validated the grouping by profiling 8,827 adipocyte nuclei by parallel scRNA-Seq (10X Chromium) sequencing(Gaublomme et al., 2019) (**Fig. 1d, Methods**), using the same RFP enrichment strategy. Similar to our first analysis, most nuclei expressed both general adipocyte and brown adipocyte markers (**Fig. S1g**), while preadipocyte markers *Cd34, Ly6a, Pdgfra* (**Fig. S1h**) and other stromal cell markers such as *Cd3, Cd14, Cd16, Cd19, Cd20, Cd56* were virtually absent (data not shown). The brown adipocytes in this larger dataset partitioned into four populations P-RT-1-4 (**Fig. 1d**). *Pde3a, Cish, Atp5e, Cyp2e1* were identified as markers of populations P-RT-1, P-RT-2, P-RT-3, P-RT-4 respectively (**Fig. 1e** and **Fig. S1i**). Besides *Cyp2e1*, P-RT-4 (256 out of 8827, violet) uniquely express *Auts2* and *Atp2b4*, mirroring the expression of the C3 cell population (**Fig. 1f**). In conclusion, both analyses identify a small cell population in murine iBAT.

In order to reveal the dynamics of brown adipocytes in response to thermoneutrality (TN) and cold exposure (CE), we profiled 8,200 adipocyte nuclei from iBAT of mice kept at 30°C for 120 days and 11,432 adipocyte nuclei from iBAT of mice exposed to 8°C for 4 days. An integrated analysis for nuclei in RT, CE and TN revealed ten subsets (**Fig. 1g,h**), with P1 and P2 derived mainly from RT and CE, P3 mainly from TN, P5 from RT, P6-P10 mainly from CE (**Fig. 1i**). Interestingly, P4 contained nuclei from all three conditions and was mainly comprised of P-RT-4, P-TN-4 and P-CE-4 (**Fig. S1j, Fig. 1i**), which was comprised of 2.45%, 3.9% and 8.3% of all cells at CE, RT and TN, respectively (**Fig. 1i**), suggesting that the number of cells from this population declines upon cold stimulation and increases in the absence of sympathetic input to brown adipose tissue. Trajectory analysis showed that the three states of cells reflect the three stimulations, while P4 cells are on the early stage of each progression (**Fig. 1j and Fig. S1k**). Clustering identified five subsets of brown adipocytes in TN condition (**Fig. S1l**), marked with *Kng2* (P-TN-1), *Ryr1* (P-TN-2), *Atp5e* (P-TN-3), *Plcb1* (P-TN-4), and *Dcn* (P-TN-5), respectively (**Fig. S1n,o**). P-TN-4 cells were similar to C3 and P-RT-4 by co-expression of markers such as *Auts2, Aldh1a1, Atp2b4* (**Fig. S1p**). In the CE condition, we identified five subsets (**Fig. S1q**), marked with *Fam13a* (P-CE-1), *Pck1* (P-CE-2), *Cyyr1* (P-CE-3), *Igf1* (P-CE-4), *Arhgap15* (P-CE-5) (**Extended Data Table 2**). Similar to the TN condition, P-CE-4 corresponded to P-TN-4, P-RT-4 and C3 (**Fig. S1t**). In the integrated analysis, P4 formed a stable cluster independent of the stimulation condition with overlapping marker genes (**Fig. 1k**).

We next tested if a similar adipocyte population is present in human brown adipose tissue, by isolating nuclei from deep neck BAT of sixteen individuals followed by snRNA-seq. Unsupervised clustering of 42,295 nuclei profiles identified 10 subsets (populations a-j) (**Fig. 1l**). Population d expressed known brown and white adipocyte markers such as *ADIPOQ, PLIN1, CIDEA, PPARGC1A* (**Fig. 1m**), suggesting that it was comprised mainly of mature fat cells. Populations b and c express high level of immune cell markers (**Fig. S1u**). A Garnett classifier trained with ADIPOQ^+^ cells as reference annotated 3,599 adipocytes mainly derived from population d (**Fig. 1n**). Sub-clustering of these adipocytes identified eight subsets (H-Ad-1-8) (**Fig. 1o**). Brown adipocyte markers *PPARGC1A* and *CIDEA* were enriched in subsets H-Ad-1 and H-Ad-3-6, indicating that these are the brown adipocytes of human adipose tissue (**Fig. 1p**). Mouse P4 specific marker genes such as *ALDH1A1, ATP2B4* and *AUTS2* were expressed in all populations (**Fig. 1p**), a P4 signature gene score (**Fig. S1v**) suggests that human brown adipocytes more closely resemble mouse P4 cells.

To further characterize P4 cells, we analyzed the *in situ* expression of *Cyp2e1*, their most prominent marker gene, and found that it is restricted to the mature adipocyte fraction both in brown and white adipose tissue (**Fig. 2a**), which is consistent with previous reports(Sebastian et al., 2011). To localize its expression within the mature adipocyte fraction, we performed immunostaining of CYP2E1 in iBAT of mice which expressed GFP under the control of *Ucp1* promoter(Rosenwald et al., 2013). A distinct population of cells was identified (**Fig. 2b**), which stained positive for CYP2E1 and GFP as a surrogate for *Ucp1* expression, validating our snRNA-seq. Most of the P4 cells were located at the edge of the brown adipose tissue depot, although some were interspersed within the depot (**Fig. 2b** and **Fig. S2a**). Interestingly, our data show that both unilocular, paucilocular as well as multilocular cells stained positive for CYP2E1 (**Fig. S2a**). Brown adipose tissue in mice has been suggested to expand by recruiting new brown adipocytes at the edges of the depot, suggesting that the newly identified cell subset P4 might be comprised of newly formed brown adipocytes, in line with the trajectory analysis (**Fig. 1j** and **Fig. S1j**). This however is at odds with our observation that this population is increased under TN condition. For humans, the endothelial niche has been demonstrated to harbor a precursor pool for brown adipocytes(Min et al., 2016; Tran et al., 2012). We did not observe any distinct localization of P4 in proximity to vascular structures, suggesting that there might be a different precursor pool, giving rise to this particular cell population.

**Figure 2.**
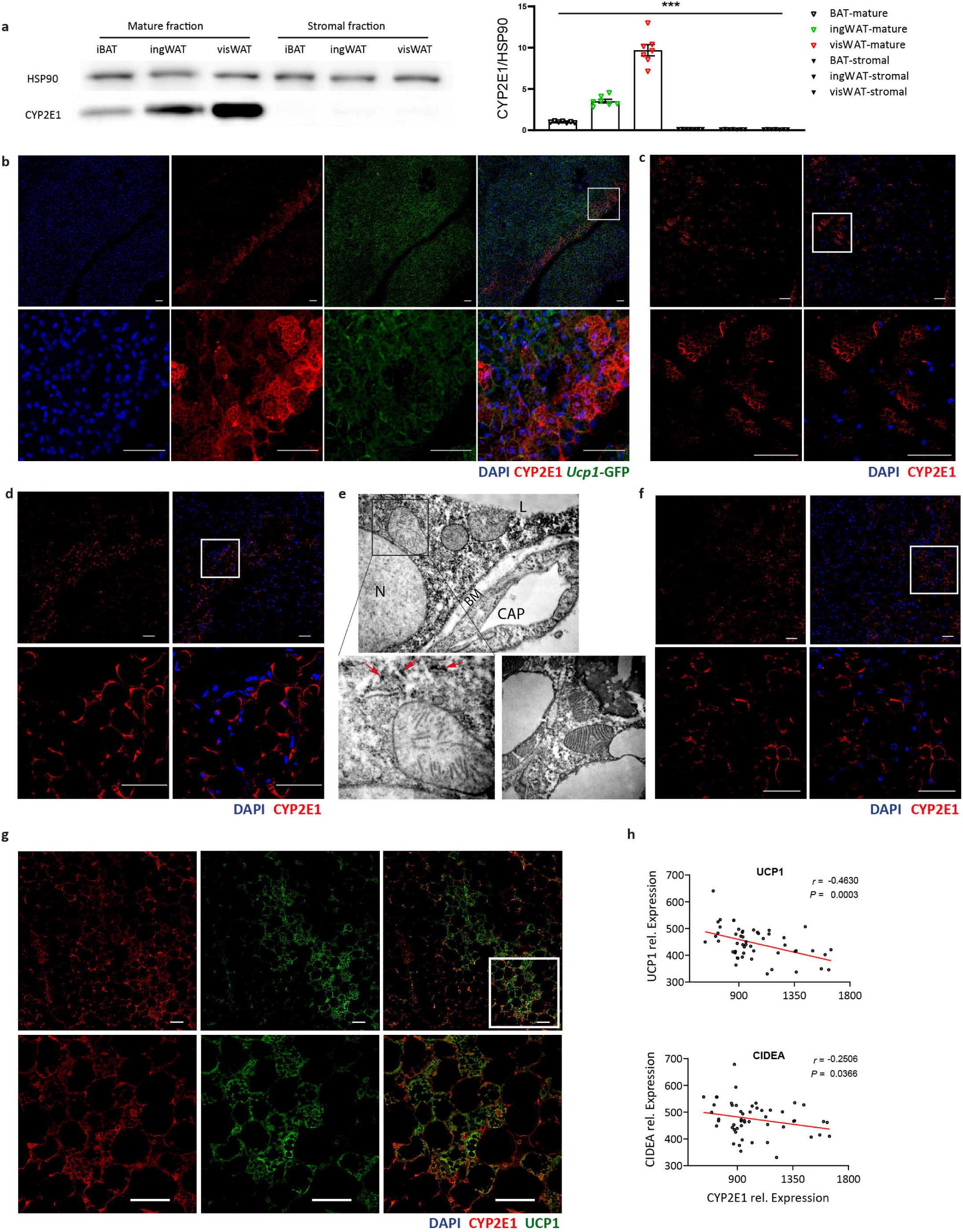
A distinct adipocyte population present in mouse iBAT, ingWAT and human deep neck BAT. (**a**) Western blot of CYP2E1 in mature adipocyte and stromal vascular fractions of three adipose tissue depots, *n* = 7, F = 167.3 DF = 41. (**b**) Immunofluorescence staining of CYP2E1 in iBAT of *Ucp1-GFP* transgenic mice at RT. (**c**) CYP2E1 immunofluorescence staining in iBAT at CE. (**d**) CYP2E1 immunofluorescence staining in iBAT at TN. (**e**) Electron microscopy analysis of CYP2E1 antibody stained brown adipose tissue, red arrows indicate CYP2E1 positive staining. Upper panel shows a cell with positive CYP2E1 staining, lower panel shows a cell devoid of CYP2E1 staining. (**f**) CYP2E1 immunofluorescence staining in ingWAT at RT. (**g**) Immunofluorescence staining of CYP2E1 and UCP1 in human deep neck brown adipose tissue. (**h**) Correlations of *CYP2E1* mRNA with *UCP1 and CIDEA* levels in human adipose tissue. Results are reported as mean ± SEM. Statistical significance was calculated using ANOVA test (**a**); *** for *P* < 0.001. Scale bar is 50 μm.

Because the snRNA-seq profiles indicated that the proportion of P4 cells may vary under different stimuli known to affect BAT function (**Fig. 1i**), we analyzed how the P4 population abundance and localization changed upon exposure to TN (120 days) or CE (4 days) (**Fig. 2c**). The number of P4 cells decreased after CE, while under TN condition P4 cell numbers increased (**Fig. S2b**), validating our observations from snRNA-seq (**Fig. 1i**). The cells remained localized within the depot under all conditions. These data suggest that sympathetic nervous system (SNS) innervation is involved in the recruitment of these cells in an inverse fashion compared to SNS mediated mature brown adipocyte recruitment(Nguyen et al., 2017). To perform a more detailed analysis of these cells we acquired electron microscopy images from CYP2E1 immunostained iBAT. In accordance with light and fluorescent microscopy data, CYP2E1-positive cells exhibited a brown adipocyte-like ultrastructure, bearing different amounts of lipid droplets. Their mitochondria were smaller and contained few and randomly oriented cristae (**Fig. 2e**) with an ultrastructure similar to those previously described in iBAT of rats. In addition, we observed a high percentage of CYP2E1^+^ P4 cells in ingWAT both at RT and after CE. At RT, CYP2E1^+^ cells were mainly unilocular (**Fig. 2f**), while after CE there were both multilocular and unilocular cells (**Fig. S2b**). In visWAT there was a large number of CYP2E1^+^ cells that had an exclusive unilocular shape (**Fig. S2c**). To assess the localization of P4 cells in human BAT, we stained CYP2E1 in deep neck adipose tissue samples obtained from patients undergoing thyroid surgery(Perdikari et al., 2018). In accordance with both *in situ* analyses of mouse tissue, both multilocular and unilocular cells stained positive for CYP2E1, suggesting that these cells constitute a population within the tissue that can acquire different morphological phenotypes (**Fig. 2g, Fig. S2d**). Consistent with the snRNA-seq data, the number of P4 like cells was substantially higher in human deep neck adipose tissue than mouse adipose tissue. This is in line with the fact that humans probably spend most of their time under conditions close to thermoneutrality(Keijer et al., 2019). Given the aberrant mitochondrial structure as well as the exclusive localization within the mature adipocyte fraction, we analyzed human adipose tissue samples from obese and overweight patients which underwent a program of weight loss(Perdikari et al., 2018). Interestingly, expression of *CYP2E1* was inversely correlated albeit weakly, with *UCP1* and *CIDEA* expression in these samples as well as with circulating glucose levels (**Fig. 2h** and **Fig. S2f**). Taking into account the altered mitochondrial structure that suggests that P4 cells have a compromised mitochondrial activity coupled to the inverse correlation with UCP1 we aimed to explore the hypothesis that P4 cells are associated with reduced BAT activity. Therefore, we examined genes selectively expressed in P4 in our snRNA-seq (**Extended Data Table 5**). Interestingly, one gene that is co-expressed exclusively with *Cyp2e1* is *Aldh1a1*, which has been implicated in adipose tissue thermogenesis(Kiefer et al., 2012). Co-staining of both markers in mouse iBAT (**Fig. 3a**) showed a complete overlap, suggesting that only P4 brown adipocytes express ALDH1A1. Moreover, ALDH1A1 is exclusively expressed in mature adipocytes in iBAT (**Fig. 3b**). The findings reporting that loss of ALDH1A1 has been associated with increased BAT functionality(Kiefer et al., 2012) together with our observation that CYP2E1/ALDH1A1^+^ cells have aberrant mitochondrial structure, supports the hypothesis that these cells are brown adipocytes with reduced functionality.

**Figure 3.**
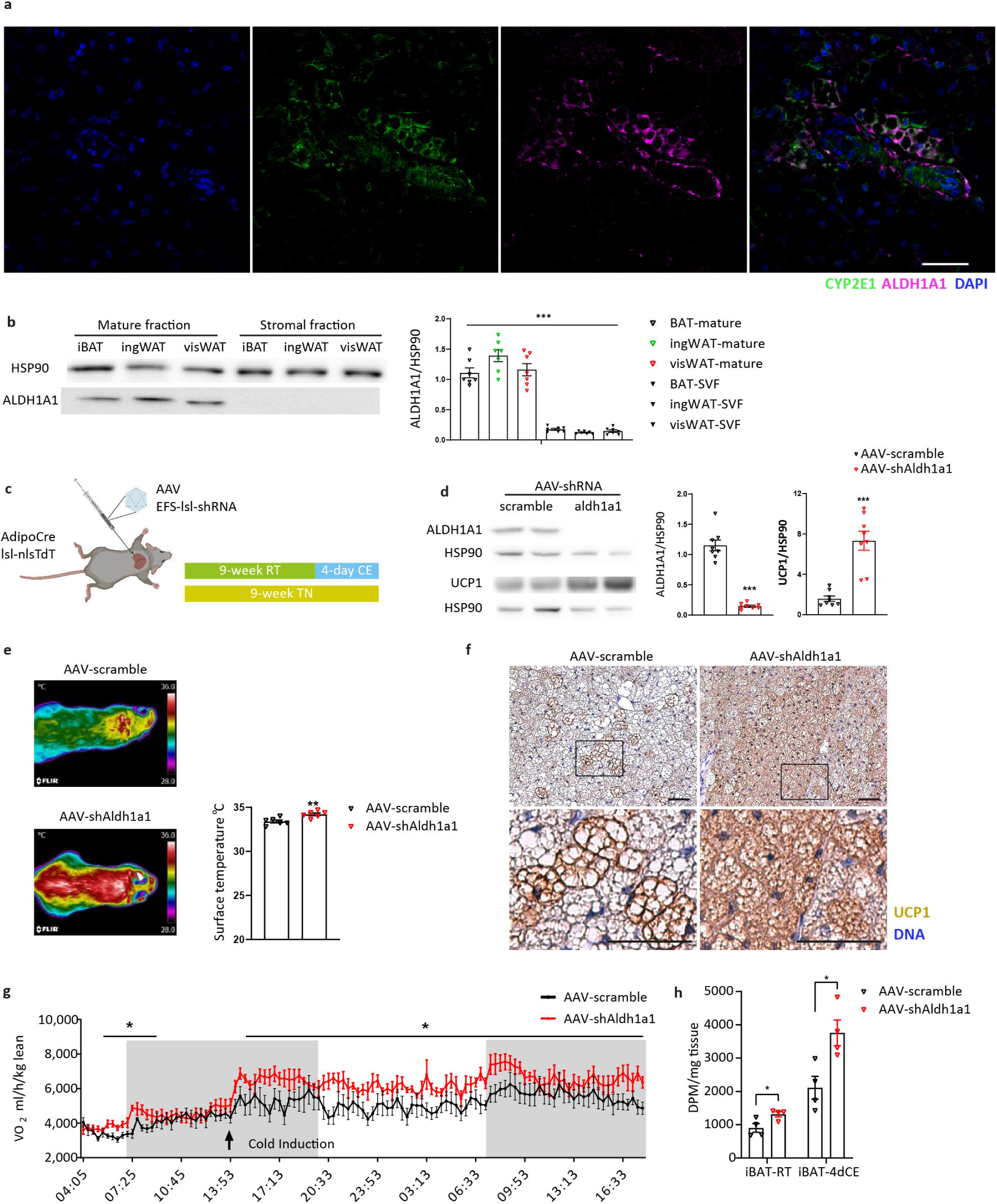
AAV-shRNA-mediated knockdown of Aldh1a1 in mature adipocytes of iBAT promotes thermogenesis and whole-body energy expenditure in mice. (**a**) Immunofluorescence co-staining of CYP2E1 and ALDH1A1 in iBAT at RT. (**b**) Western blot of ALDH1A1 in mature adipocyte and stromal vascular fractions of three adipose tissue depots, *n* = 7, F = 78.06, DF = 41. (**c**) Schematic illustration and work flow of the AAV injection experiment. (**d**) Protein levels of ALDH1A1 and UCP1 upon AAV-shRNA mediated knockdown of *Aldh1a1* in iBAT, *n* = 8, df=14, t_(ALDH1A1)_ = 11.10, t_(UCP1)_ = 5.82. (**e**) Surface temperature of AAV injected mice with 4 days of cold exposure, *n* = 6, t = 3.55, df = 10. (**f**) Immunohistochemical staining of UCP1 in iBAT from AAV injected mice at CE. (**g**) Time resolved oxygen consumption of AAV injected mice with cold induction, *n* = 5, df = 8, t_(RT)_ = 2.57, t_(CE)_ = 2.33 (**h**) Glucose uptake in iBAT of AAV injected mice, *n* = 4, df = 6, t_(RT)_ = 2.74, t_(CE)_ = 3.19. Results are reported as mean ± SEM. Statistical significance was calculated using ANOVA test (**b**); two-tailed unpaired Student’s T-test (**d** - **h**); *** for *P* < 0.001, ** for *P* < 0.01, * for *P* < 0.05. Scale bar is 50 μm.

To test this hypothesis *in vivo*, we generated Adeno Associated Viruses (AAVs) to target *Aldh1a1* expression exclusively in mature adipocytes in the iBAT depot (**Fig. S3a**). Such a system is required, since neither Cre-driver line exists, which can be used to target specifically P4 given its expression in both UCP1^+^ and UCP1^-^ cells. To test the specificity of the system we infected iBAT of AdipoCre-nucRed mice with the AAV expressing a shRNA against *Aldh1a1* both at RT with a subsequent CE as well as under TN conditions (**Fig. 3c**). We observed efficient knockdown of ALDH1A1 in iBAT (**Fig. 3d**) and exclusive targeting of mature adipocytes as evidenced by the co-expression of tdTomato and GFP (**Fig. S3b**). We did not find any changes in expression in visWAT and only a 10% decrease in ingWAT (**Fig. S3c**), which was expected given the specific Cre-expression of the chosen model(Eguchi et al., 2011). Following CE, mice with ablated *Aldh1a1* showed a significantly higher surface temperature (**Fig. 3e**) and UCP1 levels in iBAT (**Fig. 3d,f**) than control mice. This is in line with previous reports indicating that ablation of *Aldh1a1* leads to a protection from obesity concomitant with an increase in brown adipose tissue function(Kiefer et al., 2012; Ziouzenkova et al., 2007). This increase in UCP1 levels and body temperature was even more pronounced under TN conditions (**Fig. S3d**-**f**). Lastly, mice with *Aldh1a1* ablated selectively in iBAT showed higher oxygen consumption (**Fig. 3g**) and glucose uptake (**Fig. 3h**) at RT, which was even more pronounced after mice were exposed to cold. In light of the small percentage of P4 cells in iBAT (∼2.9%) this finding is of high interest as it suggests that either P4 cells following *Aldh1a1* ablation become the main contributors to systemic energy expenditure or that these cells interact with other cells to modulate their functionality.

To study this phenomenon, we cultured SVF from iBAT and ingWAT and differentiated these cells into mature adipocytes, *in vitro*. Similar to the *in vivo* data we observed a heterogeneous cell mixture in which approximately 12% of differentiated adipocytes stained positive for ALDH1A1 and CYP2E1 in iBAT derived adipocytes and 16% stained positive for both genes in ingWAT derived adipocytes (**Fig. 4a, Fig. S4a,b**). These data suggest that P4 cells arise from committed precursors, which is in accordance with a previous report demonstrating that the heterogeneity of the adipocyte precursors determines adipocyte heterogeneity to a certain extent(Wu et al., 2012). Similar to our *in vivo* results, expression of both *Cyp2e1* and *Aldh1a1* is induced upon adipocyte development similar to *Adipoq* (**Fig. 4b,c**). To modulate the function of these cells and confirm our *in vivo* findings, we ablated the expression of *Aldh1a1* by siRNA-mediated knockdown (KD), which led to an efficient repression of expression (**Fig. 4d**). SiRNA mediated KD of *Aldh1a1* did not affect the degree of adipogenesis or cell number (**Fig. 4e, Fig. S4c,d**). Ablation of *Aldh1a1*, however, led to an induction of UCP1 expression (**Fig. 4f**) and other brown adipocyte specific genes (**Fig. S4e**) most likely due to an increased number of UCP1 positive cells (**Fig. 4e**). Similar to the *in vivo* observation this increase is puzzling as only 12% of the cells in culture are positive for CYP2E1/ALDH1A1 and hints at a regulatory function of P4 in controlling tissue thermogenic capacity of other brown adipocytes. To test this hypothesis, we performed respirometric analysis of *ex vivo*-differentiated adipocytes from iBAT, in which we ablated gene expression of *Aldh1a1*. In line with the observed changes in UCP1 protein expression, oxygen consumption of these cells strongly increased (**Fig. 4g**), reminiscent of an increase in brown adipocyte functionality. Similarly, we observed an increase in ECAR suggesting an increased glycolytic flux (**Fig. S4f**).

**Figure 4.**
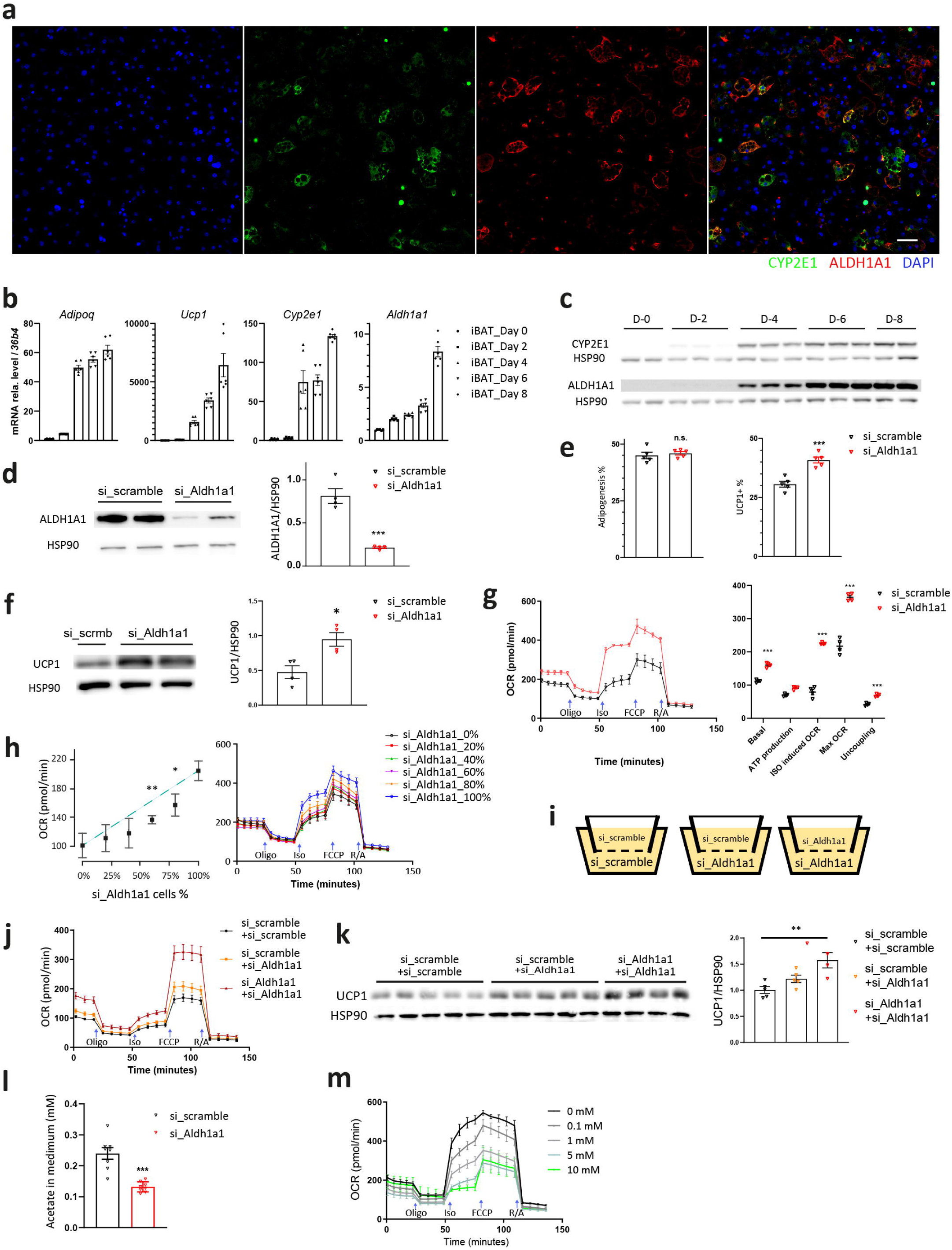
Adipocyte heterogeneity and function is conserved in ex vivo differentiated primary brown adipocytes. (**a**) Immunofluorescence staining of ALDH1A1 and CYP2E1 in *ex vivo* differentiated brown adipocytes, *n* = 6. (**b**) mRNA level of *Adipoq, Ucp1, Cyp2e1*, and *Aldh1a1* during brown adipocytes differentiation, *ex vivo.* (**c**) Protein levels of CYP2E1 and ALDH1A1 during brown adipocytes differentiation, *ex vivo*. (**d**) Protein levels of ALDH1A1 in brown adipocytes with *Aldh1a1* siRNA mediated knockdown, *n* = 4, t = 6.97, df = 6. (**e**) Percentage of differentiated and UCP1+ cells % (*n* = 5 experiments, t = 5.73, df = 8) of primary brown adipocytes after *Aldh1a1* knockdown. (**f**) Protein levels of UCP1 in brown adipocytes following *Aldh1a1* siRNA mediated knockdown, *n* = 4, t = 3.49, df = 6. (**g**) Cellular respiration in iBAT *ex vivo* differentiated cells with *Aldh1a1* siRNA mediated knockdown, *n* = 4, df = 6, t_(Basal)_ = 11.9, t_(Atp)_ = 7.16, t_(Iso)_ = 17.8, t_(Max)_ = 11.3, t_(Uncoupling)_ = 9.75, blue arrows indicate compound injection: Oligomycin 1 μM (Oligo), Isoproterenol 1μM (Iso), Carbonyl cyanide-4- (trifluoromethoxy)phenylhydrazone 1 μM (FCCP), Rotenone 3 μM and Antimycin 3 μM (R/A). (**h**) Isoproterenol induced cellular respiration in iBAT *ex vivo* differentiated cells. Cells transfected with *Aldh1a1* or scramble siRNA were mixed in different ratios, *n* = 4, df = 3, t_60%_ = 6.46, t_80%_ = 3.49. (**i**) Schematic illustration of the co-culture experiment. Cell transfected with *Aldh1a1* or scramble siRNA were cultured in the bottom or top chamber, as indicated. (**j**) Cellular respiration of co-cultured cells in the bottom well, *n* = 6. (**k**) UCP1 protein levels of co-cultured cells in the bottom well, *n* = 4-5, F = 9.24, DF = 13. (**l**) Acetate levels in culture media of *Aldh1a1* siRNA knock-down mature adipocytes, *n* = 8, t = 5.58, df = 14. (**m**) Cellular respiration in iBAT *ex vivo* differentiated cells treated with various levels of acetate during differentiation from day 2 to 8, *n* = 5. Results are reported as mean ± SEM. Statistical significance was calculated using two-tailed unpaired Student’s T-test (**d**-**g, i**) or using ANOVA test (**k**); *** for *P* < 0.001, ** for *P* < 0.01, * for *P* < 0.05. Scale bar is 50 μm.

Given the increased number of UCP1 positive cells, the increase of UCP1 protein levels coupled to the strong effect on energy metabolism when ALDH1A1 is ablated in only 12% of the cells, we speculated that the effect we observed is not cell autonomous but rather due to a paracrine interaction between P4 and other thermogenic cells in culture. To test this hypothesis, we treated *in vitro* differentiated SVF cells with *Aldh1a1* siRNA(Yang et al., 2017) or scrambled siRNA and mixed them in different ratios. We observed an *Aldh1a1* siRNA dose dependent increase of oxygen consumption and ECAR in this culture, which was substantially stronger than the expected linear increase (**Fig. 4h** and **Fig. S4g**), suggesting that P4 cells interact with other thermogenic adipocyte in culture to modulate their activity. To support this hypothesis, we performed co-culture experiments in which we cultured cells with or without ablation of *Aldh1a1* in an insert-based system (**Fig. 4i**). When we assayed the cells in the bottom chamber for oxygen consumption and ECAR, we observed that reduction of *Aldh1a1* only in the bottom chamber led to a mild increase while ablation in both chambers had a similar effect as ablation in culture (**Fig. 4j** and **Fig. S4h**). This data was in line with the observed increase in UCP1 expression only after ablation of *Alh1a1* in both chambers (**Fig. 4k**). Taken together, these data demonstrate that *Aldh1a1*/*Cyp2e1*^+^ cells express a paracrine factor, which represses thermogenic function in brown adipocytes.

*Aldh1a1* ablation has been demonstrated to lead to the increase in retinaldehyde, which activates brown adipocyte function(Kiefer et al., 2012; Yasmeen et al., 2013; Ziouzenkova et al., 2007). Based on our data we propose that in addition to this metabolite, an inhibitory factor, which affects other brown adipocytes in a paracrine fashion, exists. Previous work suggests that besides retinaldehyde, ALDH1A1 can convert acetaldehyde to acetate thereby regulating systemic function(Li et al., 2018; Liu et al., 2018; Mews et al., 2019). We therefore quantified acetate levels in the supernatant of cells in which *Aldh1a1* expression was ablated and observed a 2-fold decrease in acetate levels under these conditions (**Fig. 4l**). As demonstrated before, brown adipocytes express high levels of HMG-CoA synthase 2(Balaz et al., 2019), the rate-limiting enzyme of ketone body synthesis in addition to BDH1(Perdikari et al., 2018), which is required to convert acetoacetate to generate β-hydroxybutyrate. Under these conditions, it is possible that acetoacetate is converted into acetaldehyde, which is metabolized by ALDH1A1 to form acetate. To test whether acetate influences brown adipocyte function, we titrated brown adipocytes with acetate and measured cellular respiration. We observed a dose dependent decrease in brown adipocyte function, when acetate levels were modulated similar to the amounts observed in circulation(Müller et al., 2019) (**Fig. 4m**). Acetate has been implicated in many physiological processes, however the data in relation to obesity is conflicting(González Hernández et al., 2019). This might be due to the fact that acetate can be produced by the gut-microbiota, which can lead to changes in portal acetate concentrations as well as changes in liver function. Similarly, it is possible that acetate can act as a paracrine factor within certain tissues independent of circulating concentrations. A recent cross-sectional study revealed that circulating but not fecal acetate is negatively correlated with whole-body lipolysis and insulin sensitivity, indicating that acetate might be an important regulator of energy metabolism(Müller et al., 2019).

In conclusion, we provide here the first single cell analysis of mature brown adipocytes in both mice and humans. The RFP based nuclei selection allowed us to circumvent the need for prolonged tissue processing, digestion and adipocytes centrifugation, each of which can introduce biased cell loss. Conversely, isolation of nuclei from frozen tissue allowed us to preserve native cell populations and states. Based on these analyses we identified a rare subpopulation of mature adipocytes, which cannot be grouped into either the brown or the white cluster. This CYP2E1/ALDH1A1^+^ cell population controls the thermogenic function of other adipocytes within the depot in a paracrine fashion by modulating the short-chain fatty acid acetate levels.

## Figure legends

**Fig. S1.**
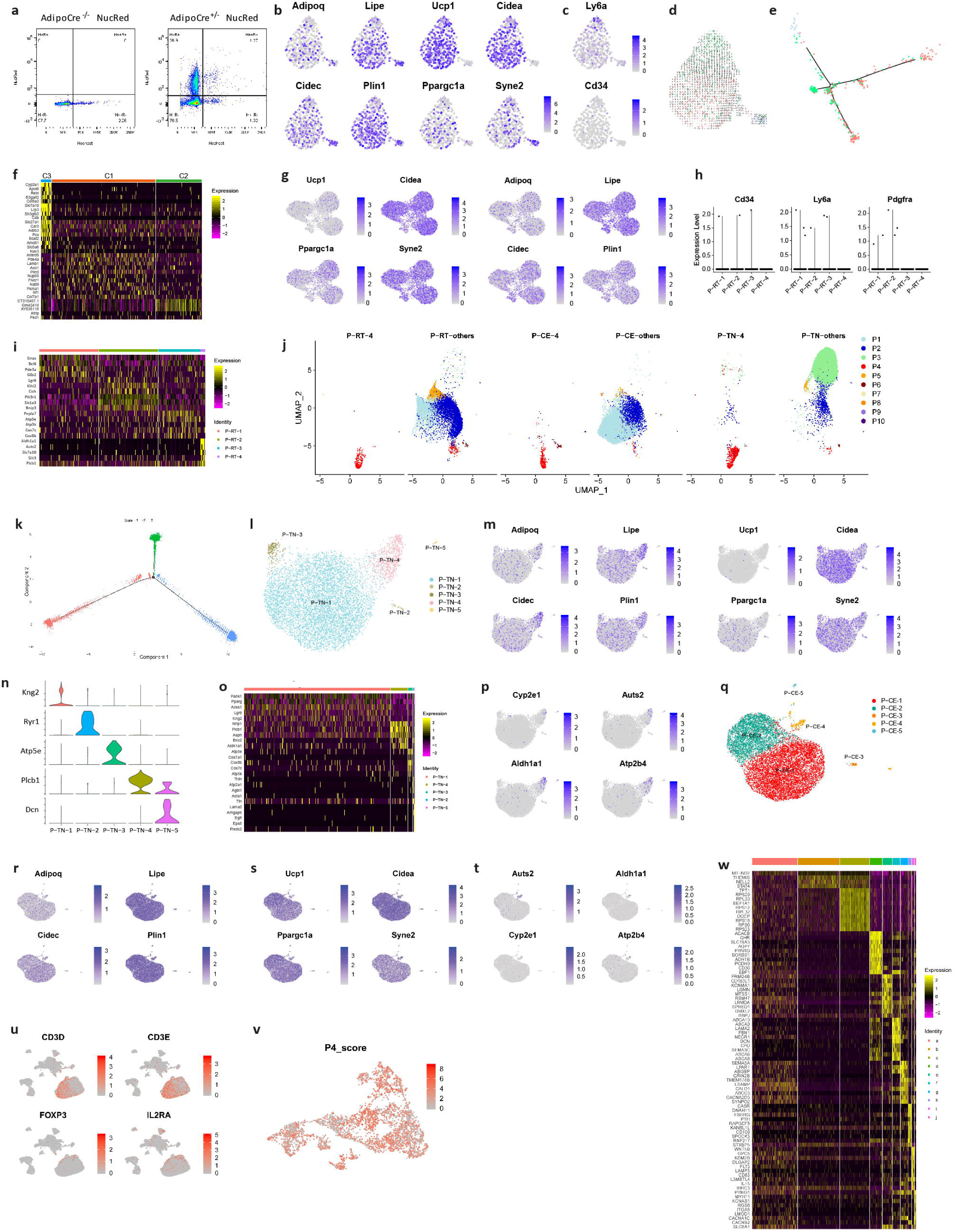
(**a**) Nuclei FACS plot of *AdipoCre* ^-/-^ *NucRed* and *AdipoCre* ^-/+^ *NucRed*. (**b**-**f**) Single-nucleus RNA sequencing of adipocytes in iBAT at RT, by Smartseq2. (**b**) Feature plots of *Adipoq, Lipe, Cidec, Plin1, Ucp1, Cidea, Ppargc1a*, and *Syne2* in brown nuclei. (**c**) Feature plot of *Cd34 and Ly6a* in brown nuclei. (**d**) RNA velocity trajectory of brown adipocytes. (**e**) Pseudotime plot of brown adipocytes using monocle (**f**) Heat map of signature genes for each population of brown nuclei. (**g-k**) Single-nucleus RNA sequencing of adipocytes in iBAT at RT, by 10x. (**g**) Feature plots of *Ucp1, Cidea, Ppargc1a, Syne2, Adipoq, Lipe, Cidec*, and *Plin1*. (**h**) Violin plot of *Cd34, Ly6a* and *Pdgfra*. (**i**) Heat map of signature genes for each population. (**i**) P-RT-4, P-CE-4 and P-TN-4 cells shown in the integrated UMAP plot. (**k**) Pseudotime plot of integrated mouse brown adipocyte snRNA seq data, grouped by states. (**l**-**p**) Single-nucleus RNA sequencing of adipocytes in iBAT at TN, by 10x. (**l**) Unsupervised clustering shown in UMAP plot. (**m**) Feature plots of *Adipoq, Lipe, Cidec, Plin1, Ucp1, Cidea, Ppargc1a* and *Syne2*. (**n**) Violin plot of *Kng2, Plcb1, Atp5e, Ryr1, Dcn*. (**o**) Heat map of signature genes for each population. (**p**) Feature plots of *Cyp2e1, Auts2, Aldh1a1*, and *Atp2b4*. (**q**-**t**) Single-nucleus RNA sequencing of adipocytes in iBAT at CE, by 10x. (**q**) Unsupervised clustering shown in UMAP plot. (**r**) Feature plots of *Adipoq, Lipe, Cidec*, and *Plin1*. (**s**) Feature plots of *Ucp1, Cidea, Ppargc1a*, and *Syne2*. (**t**) Feature plots of *Cyp2e1, Auts2, Aldh1a1*, and *Atp2b4*. (**u**) Feature plots for *CD4, PTPRC, CFD*, and *CCDC80* of snRNAseq for human deep neck brown adipose tissue. (**v**) P4 score feature plot of human adipocyte clustering, P4_score consists of the expression level of signature marker genes of P4: CYP2E1, ALDH1A1, CHST1, ATP2B4 and AUTS2. (**w**) Heat map of signature genes for each population in human brown adipose tissue.

**Fig. S2.**
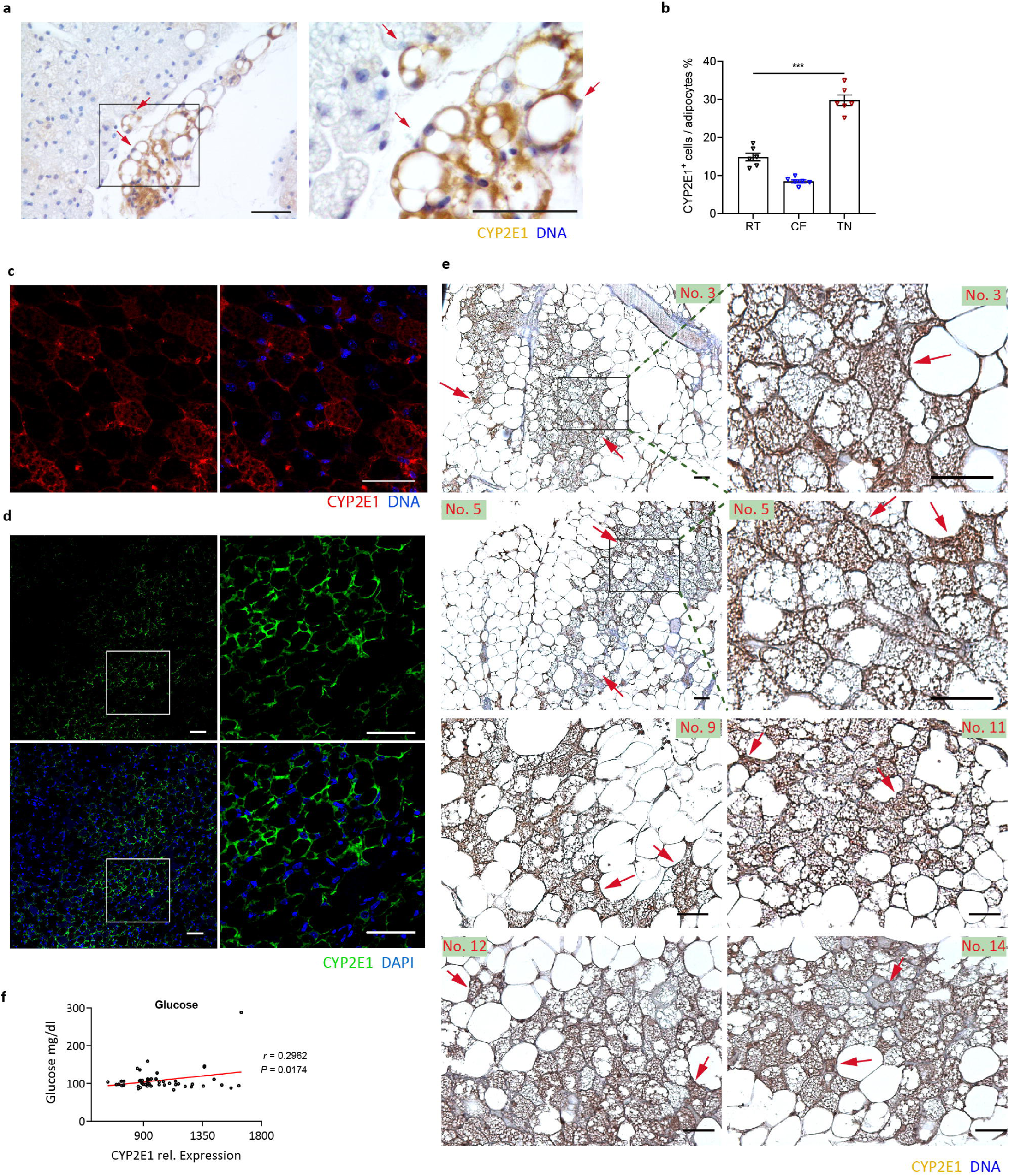
(**a**) CYP2E1 immunohistochemical staining in iBAT at RT, red arrows indicate CYP2E1 positive staining. (**b**) Quantifications of CYP2E1+ cells in iBAT at RT, CE or TN, based on images related to **Fig. 2 b-d**, n = 6, F = 112.2, DF = 17. (**c**) CYP2E1 immunofluorescence staining in ingWAT at CE. (**d**) CYP2E1 immunofluorescence staining in visWAT at RT. (**e**) Immunohistochemical staining of CYP2E1 in human deep neck brown adipose tissue, individual No.3, No.5, No. 9, No. 11, No. 12, No. 14, red arrows indicate CYP2E1 positive staining. (**f**) Correlations of *CYP2E1* mRNA with circulating glucose. Scale bar is 50 μm.

**Fig. S3.**
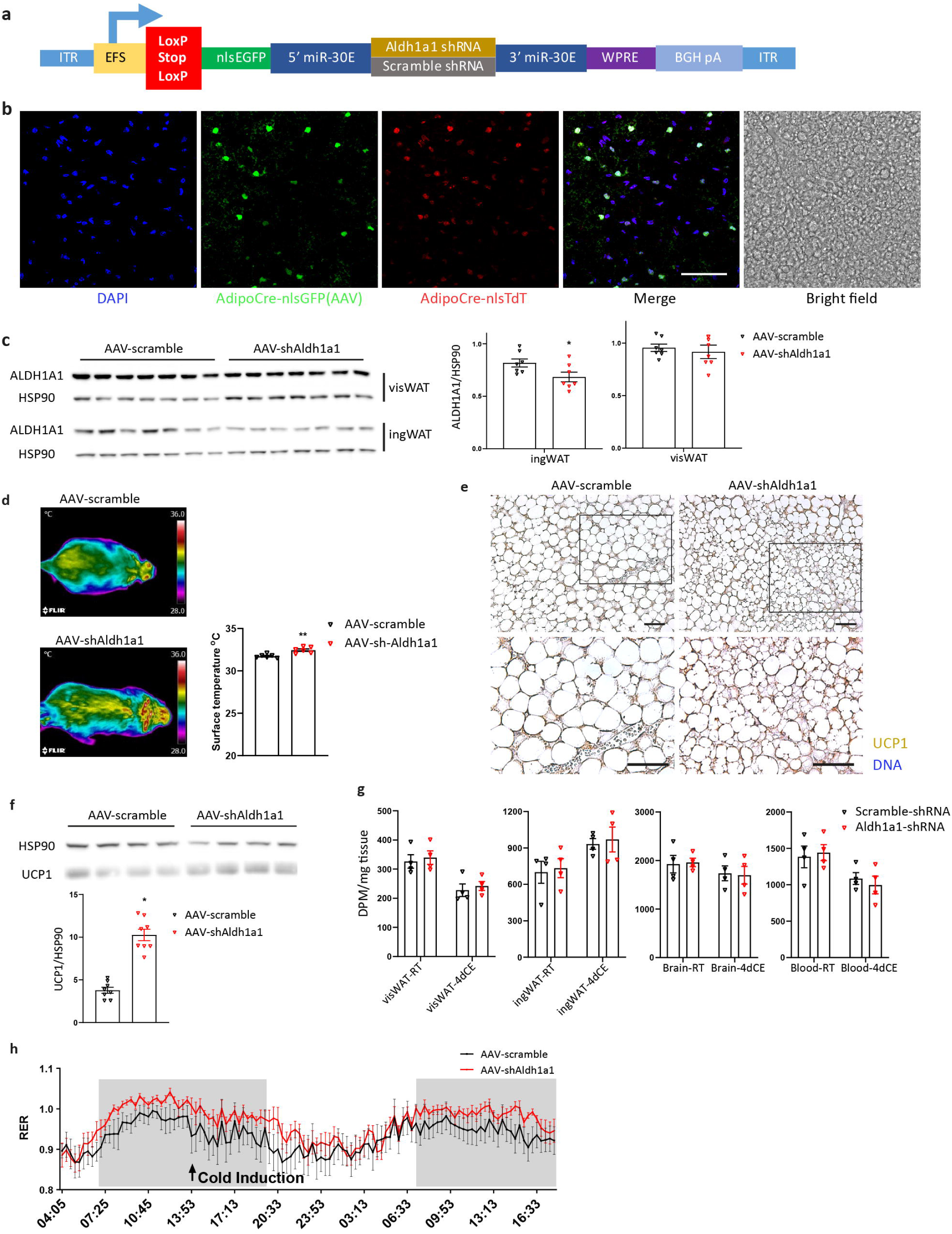
(**a**) Schematic map of the AAV construct. (**b**) Confocal images of iBAT from AAV injected mice. (**c**) Immunoblot of ALDH1A1 in ingWAT (*n* = 7, t = 2.23, df = 12) and visWAT (*n* = 7, t = 0.53, df = 12) following AAV-iBAT injection. (**d**) Surface temperature of AAV injected mice at TN, *n* = 6, t = 4.17, df = 10. (**e**) Immunohistochemical staining of UCP1 in iBAT from AAV injected mice at TN. (**f**) Protein levels of UCP1 in iBAT from AAV injected mice at TN, *n* = 8, t = 8.58, df = 14. (**g**) Glucose uptake of ingWAT, visWAT, brain, blood in mice injected with AAV, n = 4. (**h**) Time resolved respiratory exchange ratio of AAV injected mice with cold induction, *n* = 5. Results are reported as mean ± SEM. Statistical significance was calculated using two-tailed unpaired Student’s T-test. ** for *P* < 0.01, * for *P* < 0.05. Scale bar is 50 μm.

**Fig. S4.**
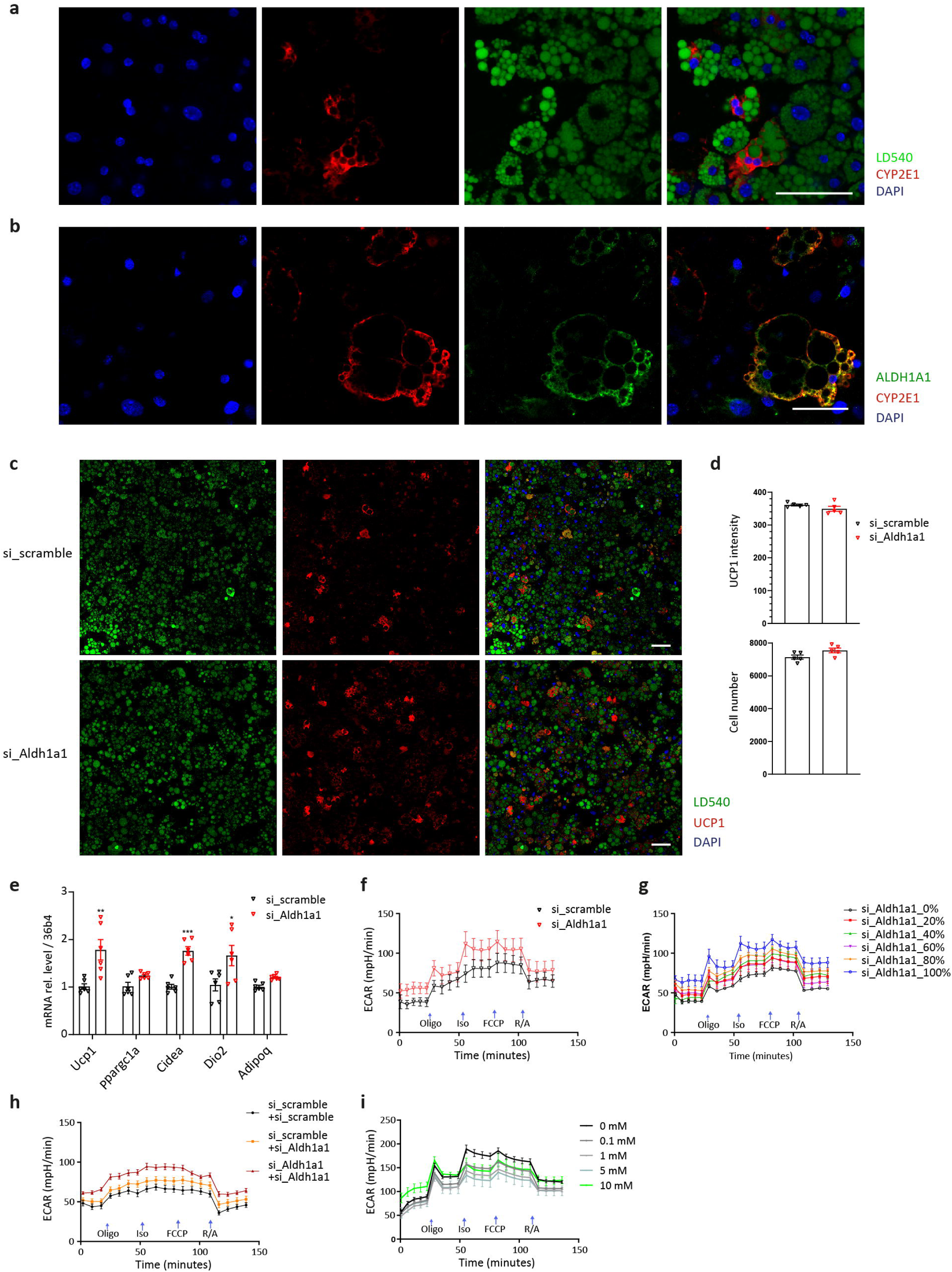
(**a**) Immunofluorescence staining of CYP2E1 and LD540 in *ex vivo* differentiated brown adipocytes. (**b**) Immunofluorescence staining of ALDH1A1 and CYP2E1 in *ex vivo* differentiated white adipocytes. (**c**) Immunofluorescence staining of UCP1 and LD540 in *ex vivo* differentiated brown adipocytes following *Aldh1a1* siRNA mediated knockdown. (**d**) Quantification of UCP1 and LD540 staining in *ex vivo* differentiated brown adipocytes following *Aldh1a1* siRNA mediated knockdown, *n* = 5. (**e**) mRNA levels of *Ucp1, Ppargc1a, Cidea, Dio2*, and *Adipoq* in *ex vivo* differentiated brown adipocytes upon *Aldh1a1* and scramble siRNA knockdown, *n* = 6, df = 10, t_(*Ucp1*)_ = 3.46, t_(*Ppargc1a*)_ = 2.59, t_(*Cidea*)_ = 7.30, t_(*Dio2*)_ = 2.49. (**f**) Time resolved ECAR in iBAT *ex vivo* differentiated cells. upon *Aldh1a1* siRNA mediated knockdown, *n* = 4. (**g**) Time resolved ECAR of iBAT *ex vivo* differentiated cells. Cells transfected with *Aldh1a1* or scramble siRNA were mixed in different ratios, *n* = 4. (**h**) Time resolved ECAR of *Aldh1a1* or scramble siRNA transfected *ex vivo* differentiated cells, which were co-cultured with *Aldh1a1* or scramble siRNA transfected primary adipocytes, *n* = 6. Schematic illustration of this experiment is shown in Fig. 4i. (**i**) Time resolved ECAR in *ex vivo* differentiated cells treated with different level of acetate during differentiation, *n* = 5. Results are reported as mean ± SEM. Statistical significance was calculated using two-tailed unpaired Student’s T-test; *** for *P* < 0.001, ** for *P* < 0.01, * for *P* < 0.05. Scale bar is 50 μm.

## Methods and Materials

### Clinical sample acquisition

The clinical study was approved by the Local Ethics Committee of the University Hospital in Bratislava, Slovakia. All study participants provided witnessed written informed consent prior to entering the study. Brown adipose tissue samples were obtained from the lower third of the neck by an experienced ENT surgeon during neck surgery under general anesthesia. The deep neck brown adipose tissue samples were taken from pre- and paravertebral space between common carotid and trachea in case of thyroid surgery and just laterally to carotid sheath in case of branchial cleft cyst surgery. In all cases, the surgical approach was sufficient to reach and sample the deep neck adipose tissue without any additional morbidity. Patients with malignant disease and subjects younger than 18 years were excluded from participation in the study. Deep neck BAT samples were collected from 16 individuals (4 male/12 female; 49.2 ± 19.0 years (22 – 77 years); BMI 24.8 ± 4.7 kg/m^2^ (16.9 – 35.2 kg/m^2^); body fat 29.1 ± 8.5 % (15.6 – 46.6 %); thyroid surgery n=14 or branchial cleft cyst surgery n=2; data are expressed as mean ± SD). Samples were cleaned immediately from blood and connective tissue, frozen in liquid nitrogen and stored at −80°C until isolation of nuclei.

### Nuclei isolation from human tissue

Nuclei were isolated following a modified nuclear isolation protocol(Drokhlyansky et al., 2019). Briefly, frozen human BAT tissues were thawed on ice, minced to 1 mm^3^ and homogenized in cold 0.1% CHAPS in Tris-HCL. The minced adipose tissue was filtered through 40 μm cell strainer, centrifuged at 500g for 5 minutes at 4 °C and the pellet was resuspended in PBS with DAPI. Nuclei suspensions were loaded to MoFlo Astrios EQ Cell Sorter and sorted into a 1.5 ml tube.

### Nuclei isolation from mouse tissue

Interscapular brown adipose tissue was harvested from seven-week-old AdipoCre-NucRed transgenic mice for each experiment. Nuclei were isolated by following a modified DroNc-seq protocol(Habib et al., 2017). Tissue was minced to 1 mm^3^ and homogenized in Nuclei EZ Lysis Buffer (#NUC101, Sigma-Aldrich) on ice and filtered through a 40 μm cell strainer. This was followed by centrifugation at 500 g for 5 minutes at 4 °C and the pellet was resuspended in PBS. Resuspended nuclei were loaded to MoFlo Astrios EQ Cell Sorter and RFP^+^ nuclei were collected individually in 384-well plate for Smartseq2 or 10x sequencing.

### Single-nucleus sequencing

Smartseq2(Chen et al., 2017) based libraries were generated following a modified Div-seq method(Habib et al., 2016). Briefly, sorted individual nuclei were reverse transcribed with oligo-dT, TSO, Super Scriptase II, RT buffer, dNTP, Betaine, MgCl_2_, RNAse inhibitor. Then the RT product was amplified by 21 PCR cycles with ISPCR primer and KAPA HiFi kit and then purified with AMPure XP beads. The PCR products were fragmented by Nextera XT kit and sequenced on Nextseq 500. Primer sequences are available upon request. 10x based libraries were generated following manufacture’s protocol. Briefly, a 1000 nuclei/ul suspension was loaded to 10x chromium with a V3 kit. Sequencing was performed using a Novaseq. Sequencing data was analyzed following a protocol based on Seurat(Satija et al., 2015) package V3.1.1 and monocle V2.14.0(Qiu et al., 2017) on R 3.6.1(Ihaka and Gentleman, 1996).

10x based libraries were acquired with the Chromium Single Cell V3.0 reagent kit following the manufacturer’s protocol (10x Genomics). Nuclei suspensions containing at ∼500 nuclei/ul were loaded into nine independent lanes. Libraries were sequenced on a Novaseq 6000 (Illumina). For data analysis, first we applied CellBender(Fleming et al., 2019) to distinguish cell-containing droplets from empty droplets; Then we applied Scrublet(Wolock et al., 2019), DoubletFinder(McGinnis et al., 2019) exclude potential doublets; meanwhile nuclei that expressed both male and female signature genes were excluded for down-stream analysis. Human Ensembl-GRCh38.p13 and mouse Ensembl-GRCm38.p5 were used for mapping. CCA^51^ from Seurat package was applied for batch correcting, clustering and signature gene identification.

### AAV production

AAV plasmid was ordered from vector builder, with shRNA sequence targeting Aldh1a1 (CCGCAATGAAGATATCTCAGAATAGTGAAGCCACAGATGTATTCTGAGATATCTTCATTG CGA) or scrambled control. 10 ug of targeting vector was co transfected with 40 ug pDP8 and 200 ul PEI (1 mg/ml) in a P15 of 293AAV cells (AAV-100, Cell biolabs), at 40% confluence. Culture medium was refreshed 24 hours post transfection. Culture medium was collected 72 hours post transfection and concentrated with AAVanced Concentration Reagent (#AAV100A-1, System Biosciences).

### AAV administration

AAV injection was performed following an established protocol^52^. AdipoCre-NucTdT mice were anesthetized with isoflurane. A longitudinal incision at interscapular region was performed to expose the brown fat depot, six injections with 10 ul of AAV (10^13^ vg/ml) were distributed in both side of brown fat.

### Primary adipocyte isolation and culture

For SVF isolation, dissected adipose tissues were minced with scissors and incubated in 1 mg/ml collagenase (#C6885-1G, Sigma-Aldrich) in collagenase buffer (25 mM NaHCO_3_, 12 mM KH_2_PO_4_, 1.2 mM MgSO_4_, 4.8 mM KCl, 120 mM NaCl, 1.4 mM CaCl_2_, 5 mM Glucose, 2.5% BSA, 1% Pen/Strep, pH=7.4) for 50 min at 37°C under agitation. Equal volume of culture media (high glucose DMEM medium (#61965026, Gibco) supplemented with 10% FBS and 1% Pen/Strep) was added and samples were centrifuged for 5 min at 300 g. The SVF pellet was resuspended in 2 ml erythrocyte lysis buffer (154 mM NH_4_Cl, 10 mM KHCO_3_, 0.1 mM EDTA, pH 7.4) and incubated for 4 min in room temperature. Samples were diluted with 10 ml culture media and filtered through 40 µm cell strainers. After 5 min of centrifugation at 300g, the supernatant was removed and the pellets were resuspended in culture media. SVF cells were seeded into a plate pre coated with collagen I (1:500, #C3867, Sigma-Aldrich) and differentiated as described previously^53^. 48h post differentiation induction, cells were cultured with maintenance cocktail (1μM rosiglitazone and 0.12μg/ml insulin) and refreshed every 48 hours.

To test the effects of acetate on brown adipocyte function, culture media was supplement with different amounts of acetate during differentiation. To quantify acetate, primary brown adipocytes on differentiation day 8 were washed three times with PBS, and incubated with maintenance cocktail for 24 hours, 100 ul culture media was collected and acetate level was quantified with Acetate Colorimetric Assay Kit (#MAK086-1KT, SIGMA) following the manufacturer’s protocol.

### Co-culture experiment

On differentiation day 4, primary cells were reverse-transfected with a pool of 3 siRNA probes. Briefly, 75,000 cells/cm^2^ were seeded into transwell inserts or receiver plates with 100nM of corresponding siRNA, which dissolved in 1.5% Lipofectamine RNAiMAX (#13778150, Invitrogen) in Opti-MEM I reduced serum medium (#31985062, Invitrogen). 48h after transfection, the inserts and receiver plates were washed with warm PBS twice, and co-cultured as described in Fig 4**i** in maintenance cocktail. Following by 4 days of co-culture, cells were collected for protein extraction or reseeded in seahorse plates at a density of 8000 cells/well for extracellular respiration experiment.

### siRNA knock down titration experiment

On differentiation day 3, primary cells were reverse-transfected with corresponding siRNA following the protocol described above. 72h post reverse-transfection, primary cells were collected for protein extraction. The scramble siRNA transfected cells were mixed at different ratios with *Aldh1a1* siRNA transfected cells (ranging from 0 to 100%) at a density of 8000 cells/well in seahorse plates for extracellular respiration experiment.

### Indirect calorimetry

Indirect calorimetry measurements were performed with the Phenomaster (TSE Systems) according to the manufacturer’s guidelines and protocols. Animals were single caged and acclimated to the metabolic cages for 48 hours before metabolic recording^54^.

### Surface temperature measurement

Surface temperature was recorded with an infrared camera at room temperature (E60; FLIR; West Malling, Kent, UK) and analyzed with FLIR-Tools-Software (FLIR; West Malling, Kent, UK).

### Radio labeled glucose tracing

Tissue radiolabeled glucose uptake was measured, as described previously^43^. Animals were fasted for 4h, then ^14^C-2-deoxyglucose at 8 mM, 14.8 MBq/kg body weight was injected by tail vein. 30 minutes after injection, blood samples were collected. Tissue was harvested, weighed and lysed in 10 volumes of 0.5 M NaOH. Radioactivity was measured by liquid scintillation counting (100 μl of lysate in 3.9 ml of Emulsifier-Safe, Perkin Elmer).

### Analysis of adipocyte differentiation

Differentiated adipocytes at day 8 were used for differentiation analysis. Briefly, cells in 96 well optical plate were washed with PBS twice and fixed with 5% formaldehyde at 4 °C for 10 min, followed by 3 times washing with PBS. Cells were stained with LD540 (100 ng/μl) for lipid droplets and Hoechst No. 33342 (100 ng/μl). For UCP1 staining, lipids were depleted by 5% acetic acid in ethanol for 10 min at − 20 °C, washed with PBS twice at RT and blocked in 0.05% triton, 5% BSA, PBS. Cells were incubated with UCP1 antibody (1:500, #ab10983, Abcam) overnight, washed twice in PBS, incubated with Alexa Fluor 488 anti-rabbit (1:500, #A-11034, Thermo) secondary antibody and DAPI, followed by three washing steps. 29 images per well were acquired with an automated microscope imaging system (Operetta, Perkin Elmer). Images were analyzed using the Operetta imaging software Harmony, as described previously.

### Histology and image analysis

Adipose tissues were excised, fixed in fresh 4% paraformaldehyde in PBS (Gibco; pH 7.4) for 24 h at 4°C, dehydrated and then embedded with paraffin. 4-micron paraffin sections were subjected to histological staining. Heat induced antigen retrieval was applied on rehydrated paraffin sections. After blocking with 5% BSA for one hour, primary antibody (1:200 UCP1, # PA1-24894, Thermo Fisher) diluted in 5% BSA was applied to sections overnight at 4 °C. After washing with PBS, a secondary antibody (Signal Stain Boost IHC, #8114, Cell Signaling Technology) was applied and the sections were washed 3 times and were detected using the DAB method (#80259P, Cell Signaling Technology). Standard hematoxylin and eosin staining was performed on rehydrated fat paraffin sections. Slides were dehydrated and covered with coverslip by resin-based mounting. All images were acquired by Axioscope A.1 (Zeiss).

### Fluorescence immunostaining of adipose cryosections

Adipose tissues from mice were excised and fixed in fresh 4% paraformaldehyde (Sigma-Aldrich) in PBS (Gibco) at pH 7.4 for 2 h at 4 °C, washed four times in PBS and cryopreserved for 30 h in 30% sucrose in PBS with stirring at 4 °C. The samples were flash-frozen on dry ice and stored at −80 °C. Brown adipose tissues were cut at −25 °C on an HM 500 O microtome (Microm) at 20 μm thickness, mounted on Superfrost plus slides (Medite) and thawed at 4 °C, blocked with 10% donkey serum in PBS for 1 h, followed by UCP1 (#ab10983, Abcam), CYP2E1(#ab28146, Abcam), ALDH1A1(#ab9883, Abcam) antibody overnight in 10% donkey serum in PBS. Sections were washed 3 times with PBS at RT, stained with Alexa 488 anti-rabbit (#A-11034, Thermo), Alexa 594 anti-goat (#A-11058, Thermo) secondary antibody and 300 nM DAPI for 1 h. Slides were embedded in ProLong® Diamond Antifade Mountant (# P36965, Themo Fisher). Native Ucp1-GFP signal was acquired without antibody staining. Fluorescence micrographs were acquired on an SP8 confocal microscope (Leica). Background was adjusted using samples without primary antibody.

### Immunoelectron microscopy

Immunoelectron microscopy was performed on fixed samples following a modified version of this protocol(Cinti et al., 1989). Briefly, after overnight fixation at 4°C, samples were reduced in small thin slices (∼1 mm X 4 mm), placed in 0.5 ml tubes and washed with PB for 5 min at room temperature; 3% H2O2 (in PBS; 5 min) was then used to block endogenous peroxidase; samples were rinsed with PBS. They were then incubated with the primary rabbit polyclonal anti-Cytochrome P450 2E1 antibody (1:50 v/v) in PBS overnight at 4°C. After a thorough rinse in PBS, samples fragments were incubated for one hour in a 1:100 v/v secondary antibody (Vector Laboratories) solution at room temperature. Histochemical reactions were performed using Vectastain ABC kit (one hour at room temperature; Vector Laboratories) and Sigma Fast 3,3′-diaminobenzidine (10 minutes; Sigma-Aldrich) as the substrate. Samples were then reduced in ∼1mm x 1mm fragments and fixed again in 2% glutaraldheyde-2% paraformaldehyde in phosphate buffer overnight at 4°C. Samples were then post-fixed in 1% osmium tetroxide, dehydrated in a graded series of acetone, and embedded in an Epon-Araldite mixture. To determine the region of interest, semi-thin sections were cut and stained with toluidine blue. Thin sections were obtained with an MT-X Ultratome (RMC, Tucson, AZ), stained with lead citrate, and examined with a CM10 transmission electron microscope (Philips, Eindhoven, The Netherlands).

### Extracellular respiration

Primary brown adipocytes were counted and plated at a density of 8,000 cells/well in a seahorse plate and cultured with cocktail described above. At differentiation day 8, mature brown adipocytes were loaded to XF_96_ Extracellular Flux Analyzer (Agilent). Mitochondrial respiration and ECAR was quantified using the Mito-stress test protocol. After measurement of basal respiration, oligomycin (1 μg/ml inhibitor of complex V) was injected to block respiration coupled to ATP synthesis. Decrease in oxygen consumption rate (OCR) following oligomycin injection reflects contribution of coupled respiration to the basal mitochondrial OCR. Uncoupled respiration was in the next step induced with isoproterenol (1 μM), to quantify the capacity of cells to dissipate energy through uncoupled respiration. FCCP (1 μg/ml), was injected to fully uncouple the mitochondrial membrane and to quantify the maximal respiratory capacity of brown adipocytes. In the last step, Rotenone (3 μM) and Antimycin A (2 μg/ml) were injected to block mitochondrial respiration (complex I and III) and estimate contribution of non-mitochondrial respiration to the measured OCR. Non-mitochondrial respiration was subtracted to obtain basal, basal uncoupled, isoproterenol-stimulated uncoupled and maximal mitochondrial respiration(Sun et al., 2018).

### Data repository

All RNA sequencing data is deposited in ArrayExpress: E-MTAB-8561 for single-nucleus RNAseq of mouse interscapular brown adipocytes at RT by SMARTseq2; E-MTAB-8562 for single-nucleus RNAseq of mouse interscapular brown adipocytes at RT, CE, TN by 10x; E-MTAB-8564 for single-nucleus RNAseq of human BAT cells.

### Western Blot

Protein samples were isolated from adipose tissue with RIPA buffer (50 mM Tris-HCl pH (7.5), 150 mM NaCl, 1mM EDTA, 1% Triton X-100, 0.1% SDS, 10% glycerol) supplemented with protease inhibitor cocktail (#11697498001, Sigma-Aldrich) and Halt Phosphatase Inhibitor (#78420, Thermo Fisher). Homogenized protein lysates were obtained by rotating at 4 °C for 30 min, followed by centrifugation at 14,000 rpm for 30 min. Protein amounts were quantified using the DC Protein Assay (Bio-Rad). For immunoblotting, protein samples were separated by SDS-PAGE on 12% polyacrylamide gels and transferred onto nitrocellulose membrane. Membranes were probed using the indicated antibodies and chemiluminescent signals was detected by a LAS 4000 mini Image Quant system (GE Healthcare). Band intensity was quantified using ImageJ. UCP1 (#ab10983, Abcam), CYP2E1(#ab28146, Abcam), ALDH1A1(#ab52492, Abcam), HSP90 (#4887, Cell Signaling Technology), HRP anti rabbit (Calbiochem).

### Oligonucleotides

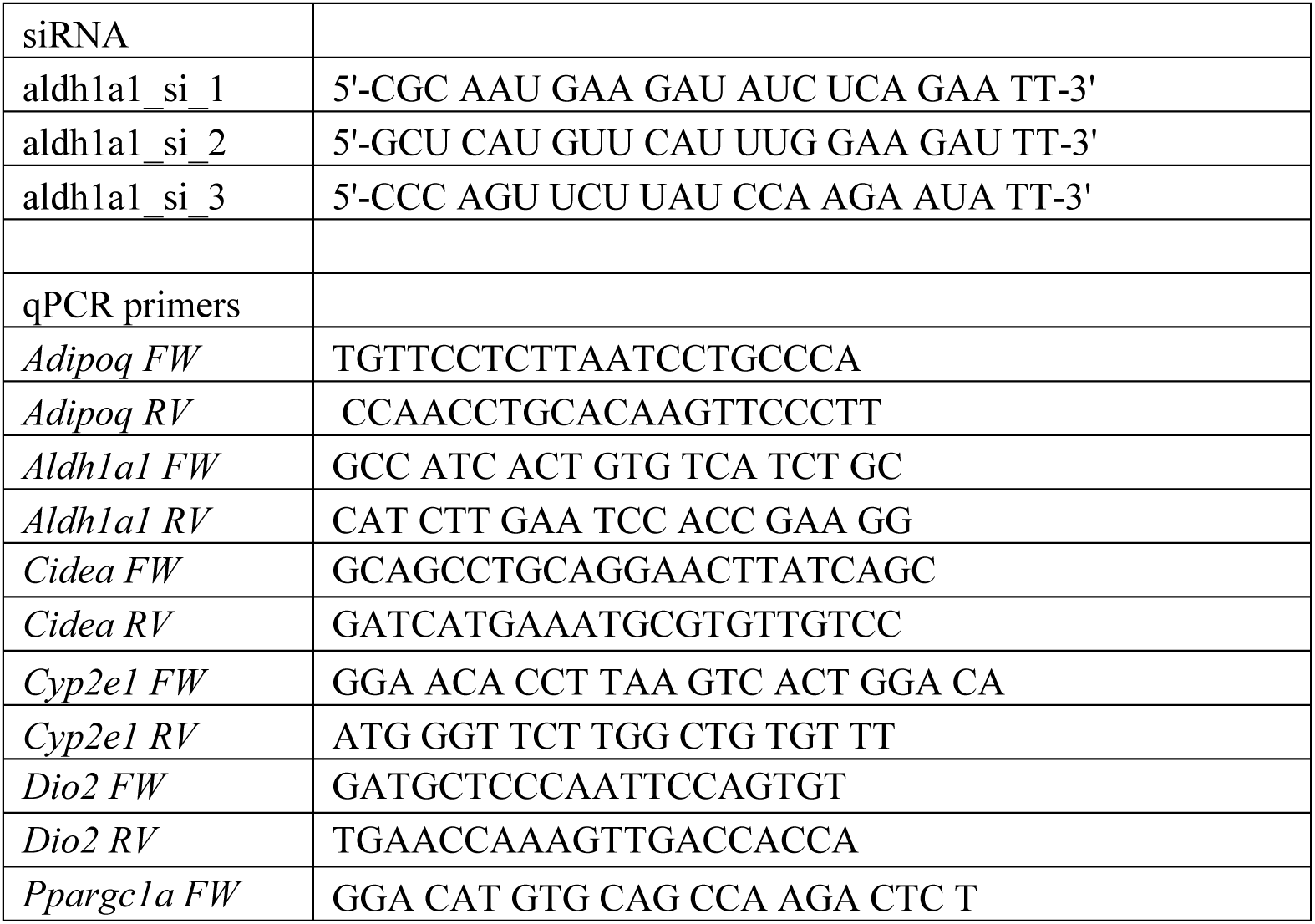
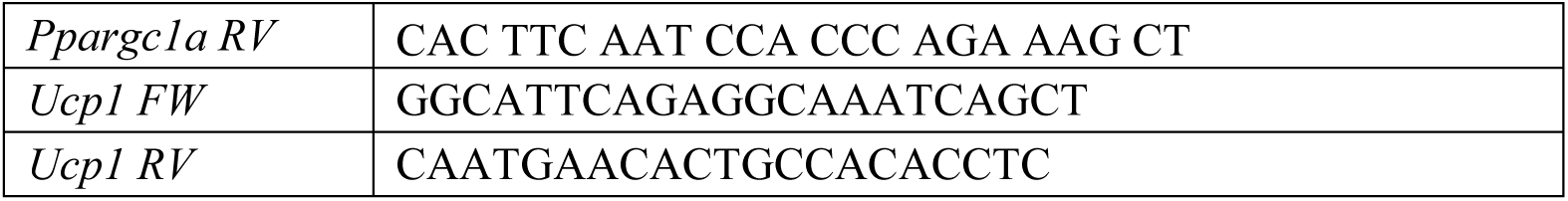

## Acknowledgements

We are grateful to Remo Freimann for assistance with nuclei FACS; Emilio Yángüez for assist with 10x and smartseq2 experiments; Evan D. Rosen for discussion and suggestions; Orr Ashenberg, Yuliang He, Umesh Ghoshdastider, Ge Tan, Bart Deplancke, Pernille Rainer and Tongtong Wang for comments on the bioinformatics analyses; Willem Koppenol and Elke Kiehlmann for histology tissue section. Data produced and analyzed in this paper were generated in collaboration with the Functional Genomics Center Zurich, Cytometry Facility of University of Zurich and the Scientific Center for Optical and Electron Microscopy of ETH. The work was supported by the Swiss National Science Foundation (SNSF 185011 to C.W.)

## Author information

W.S. conceived the study; W.S. and C.W. designed the study; W.S. and H.D. performed all of the experimental work, except those described below; W.S. analyzed the transcriptome data with input from A.R.; W.S., H.D., M.B., Z.K., and J.U. collected BAT from patients; P.S. performed surgery for human BAT collection; W.S., H.D., M.S. and E.D. developed the nuclei acquisition methods; G.C. and S.C. acquired immunoelectron microscope pictures; G.R. acquired Optifast clinical data, W.S., A.R., and C.W. wrote the manuscript; H.D., M.B., S.C., L.D., helped with editing of the manuscript.

## Contact for reagent and resource sharing

Supplementary information is available for this paper. Correspondence and requests for materials should be addressed to the lead contact Christian Wolfrum, for bioinformatic information should be directed to Wenfei Sun.

